# Wobble Vaccines: Complex Vaccine Antigen Pools Promote Increased Antibody Breadth and Cross-Strain Viral Targeting in SARS-CoV-2

**DOI:** 10.64898/2026.07.01.735277

**Authors:** Peter R. McIlroy, Wendy M. Zinzow-Kramer, Madison L. Ellis, Eduard Melief, Mohammad Ali, Hannah E. Peck, Loren E. Sasser, Daryll Vanover, Philip J. Santangelo, Mehul S. Suthar, Emily A. Voigt, Matthew C. Woodruff

## Abstract

Vaccination remains the most successful preventative measure against viral infection, but methods to stably deter rapidly-evolving pathogens have remained elusive. Vaccines capable of incorporating and anticipating viral evolution could address current challenges in seasonal vaccination efforts against SARS-CoV-2 and influenza where economic and disease burdens remain high despite decades of combined study. Rare epitope suppression (RES) is an underutilized concept within vaccine design, where humoral epitope targeting can be molded using complex antigen pools. Based in mRNA vaccine technology, ‘wobble vaccines’ represent the novel application of RES to human pathogens designed to anticipate and resist viral evolution. To establish this platform, public SARS-CoV-2 sequencing data was compiled from the first two years of the COVID-19 pandemic to identify high-diversity sites across the receptor binding domain (RBD) of the spike protein. Wobble RBD (WobbRBD) libraries reflecting that entropy were synthesized and incorporated into established self-amplifying (SA) vaccine constructs. Animals immunized with these complex antigen pools showed no obvious adverse effects. By three days-post vaccination, WobbRBD stimulated robust primary immune activation with distinctive characteristics compared to traditional single-strain vaccine modalities. By day 14, germinal centers, class switching, and antibody-secreting cells were induced, creating potent SARS-CoV-2 spike-binding IgG antibodies. Despite similar overall activation profiles, WobbRBD generated significantly increased breadth against SARS-CoV-2 variant spikes in comparison to single-strain controls – even against future-emerging strains. Taken together, wobble vaccines represent a novel method for anticipating and preventing viral escape with promising applications in SARS-CoV-2, influenza, HIV, and beyond.

## Introduction

The patient-specific diversity of the naïve B cell repertoire, combined with the stochastic cellular dynamics governing B cell selection, enables recognition of a vast array of theoretical epitopes^1, 2^. These features also result in unpredictable and unreliable outcomes in vaccination settings without direct manipulation^3, 4^. Host intrinsic factors, such as the frequency and antigen affinity of B cell precursors, result in biased inclusion and selection of activated B cells; for example, abundant or higher-affinity clones are more likely to become antibody-secreting cells (ASCs)^3, 5^. The amount of vaccine-delivered antigen and its relative avidity similarly impact the potency of both the initial B cell receptor (BCR) stimulation and the amount of costimulatory signals B cells receive, thereby influencing overall B cell selection^3, 6^. The necessity for manipulation of B cell responses is especially relevant to mutation-prone pathogens (e.g. influenza A, human immunodeficiency virus (HIV), SARS-CoV-2), where vaccines fail to promote broad and lasting protective responses as pathogens escape suboptimal immune pressures. These initial antibody responses are especially important because they establish a further imprint in both current and future B cell selection through preferential memory responses and epitope masking^7-9^. Facing these complex challenges, it is crucial to identify effective methods of creating proactive vaccine constructs to combat mutation-prone pathogens.

One extrinsic determinant of B cell selection currently underutilized in vaccine design is the relative access to particular epitopes within an antigen. Multivalent influenza vaccines, for example, introduce the hemagglutinin of multiple strains simultaneously, resulting in promotion of cross-reactive antibodies^10^. The underlying mechanism of this shift is hypothesized to be rare epitope suppression (RES). Prior work from our group has shown that as a particular epitope becomes rare in a complex antigen pool, corresponding B cell responders become less competitive for the T cell help required for ASC differentiation^11^. Thus, RES promotes production of antibodies against epitopes shared across the complex antigen pool by effectively removing antibodies targeting rare epitopes from the final humoral response.

Here, we present wobble vaccines as a novel application of RES and demonstrate initial efficacy in manipulating the B cell response to the SARS-CoV-2 spike protein receptor-binding domain (RBD). Wobble vaccines incorporate the mutational history of a pathogen to ‘wobble’ frequently-mutated sites, thereby increasing the rarity of these epitopes in the overall antigen pool, decreasing their immunodominance, and driving a relative increase in antibody titer toward conserved epitopes. This new platform displays evidence of feasibility in both design and manufacture, reasonable safety profiles, and a unique capacity to modify epitope selection in the context of vaccination. The introduction of a heterogeneous antigen pool through the RBD wobble vaccine (WobbRBD) creates a few unique changes to the resulting draining lymph node immune response without negatively impacting the overall response. Additionally, clear signs of epitope selection modification are induced by WobbRBD to increase humoral response breadth across a panel of SARS-CoV-2 variants, with evidence in favor of continued efficacy beyond the initial design timeline. With strong preliminary efficacy, wobble vaccines offer a promising and effective method of applying prior viral knowledge to creating long-lived vaccines.

## Results

### WobbRBD Library Design and Production

Previously, studies with model antigens have shown that RES is an effective method of modifying B cell epitope selection^11^. In a model vaccination system of a carrier protein conjugated to a fluorescent tag, a standard antibody response is mounted against both the carrier and tag (Fig. 1A); however, the tag response is eliminated when the carrier-tag complex is diluted by unconjugated carrier, due to competition for shared T cell help (Fig. 1B)^11, 12^. Applying this concept to vaccination, wobble vaccines are intended to combat pathogens that resist standard vaccines, where standard antibody responses target mutation-prone epitopes (e.g. influenza A hemagglutinin (HA) globular head) while conserved epitopes are ignored (e.g. HA stem) (Fig. 1C). The initial vaccine response, then, is not effective at binding or combatting future-emerging strains of this pathogen. As outlined in Fig. 1D, wobble vaccines incorporate the past mutational history of said pathogen to produce a complex pool of antigen that contains each mutation in equal proportion, thus rarifying these easily-mutated epitopes in comparison to conserved sites. RES pushes for conserved regions to be targeted, resulting in a humoral response capable of binding past, present, and future strains of this example pathogen using this knowledge of mutational probability across the protein.

**Figure 1.**
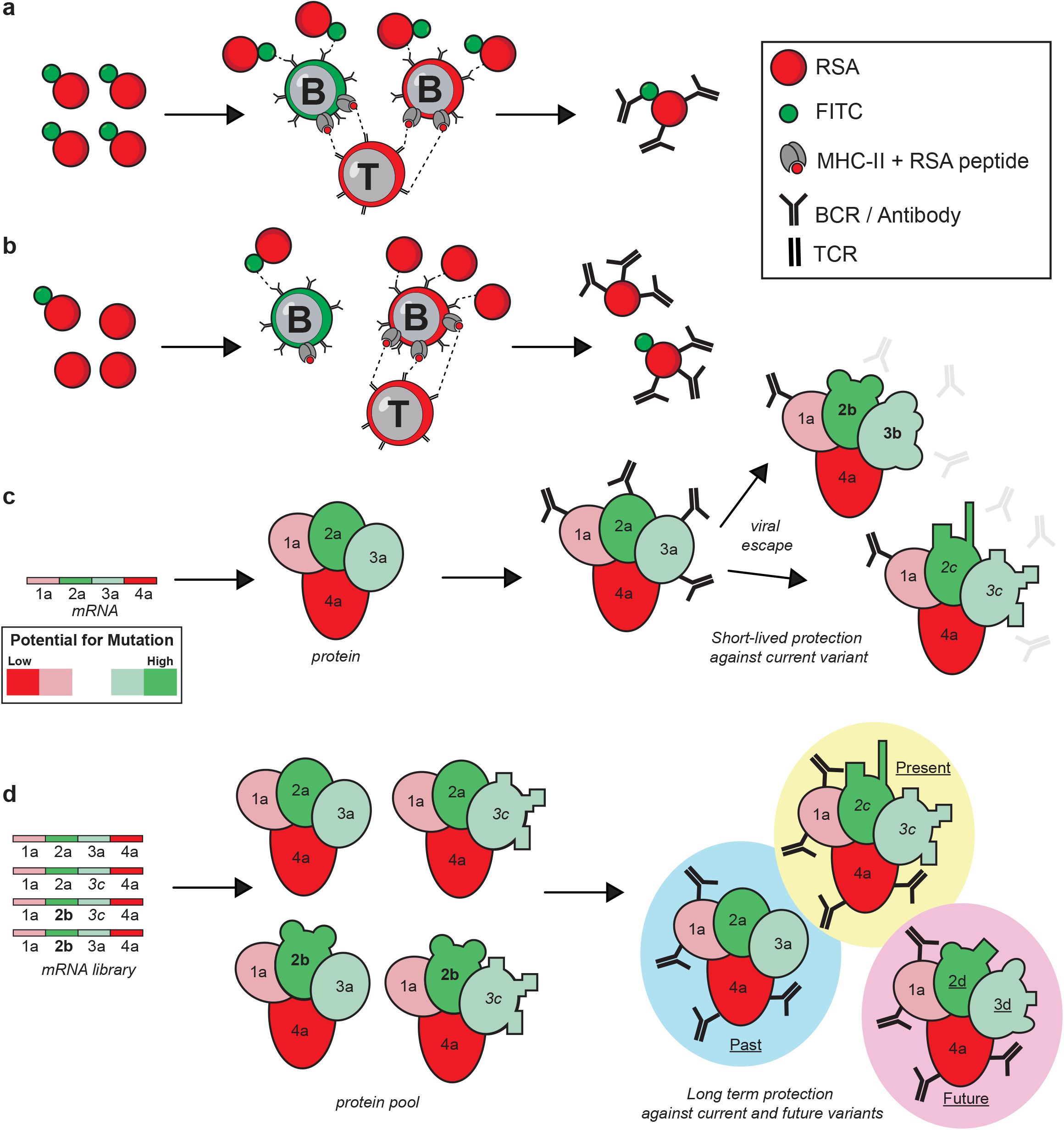
Rare epitope suppression and wobble vaccines. An illustration of the underlying mechanisms of rare epitope suppression through prior work with the carrier-tag RSA-FITC model and a hypothetical pathogenic spike protein. A) Vaccination with RSA (red) conjugated to FITC (green), leading to equal-weighted competition for T cell help to produce antibodies that bind both RSA and FITC. B) Rare epitope suppression occurs when RSA-FITC is diluted 1:3 by unconjugated RSA, leading to favored T cell help to RSA-binding B cells with increased peptide-MHC-II on their surface, and resulting in production of only RSA-binding antibodies. C) A hypothetical single strain vaccine, where regions with high potential for mutation are targeted by antibodies, leading to a loss of protection after viral escape occurs. D) A proposed schematic of a wobble vaccine response, where known heterogeneity is included in the vaccine to make these highly mutable sites ‘rare’ and thus exclude them from consideration in the final antibody response. This leads to production of antibodies that target conserved regions, leading to increased breadth and protection against past, current, and future pathogenic variants.

To test this hypothesis, wobble vaccine constructs were designed using the dataset of SARS-CoV-2 spike sequences that were captured from December 2019 through January 2022 through the Nextstrain ncov GISAID global dataset^13, 14^ (Fig. 2A). Using this snapshot of viral diversity, normalized Shannon entropy at each amino acid site within the RBD was analyzed across 10,000 sequences. Sites with a high propensity for mutation were identified with an inclusive entropic cut-off of 0.02 (Fig. 2B). This entropic cut-off included all 16 RBD mutations prevalent in variants of concern (VOC) within the assessment window, as well as an additional seven lower-entropy sites (Fig. S1A). One position with entropy above 0.02 (position 490) was excluded from library design considerations due to manufacturing constraints. DNA flex libraries containing the above identified mutations were constructed out of distinct regions of heterogeneity, here designated as wobble regions 1-6, with constant regions between them derived from the original Wuhan 2019 (WA.1) strain for final construct ligation (Table S1). Each high entropy position was mutated independently, allowing for a maximum of 8.55*10^15 possible DNA species and 1.15*10^9 possible amino acid combinations. From the identified set of amino acids that could be encoded at each wobble position, a set of codons was identified to allow for minimal point mutations from the WA.1 reference sequence while encoding the most common amino acid variants identified within the viral sequencing data. Manufactured flex libraries were verified by next-generation sequencing and computationally validated to ensure expected identity compositions at each flex locus (Fig. 2C). One WobbRBD library was manufactured for each of the six wobble regions (WobbRBD 1-6), with a final combined library containing all 6 regions (WobbRBD 7).

**Figure 2.**
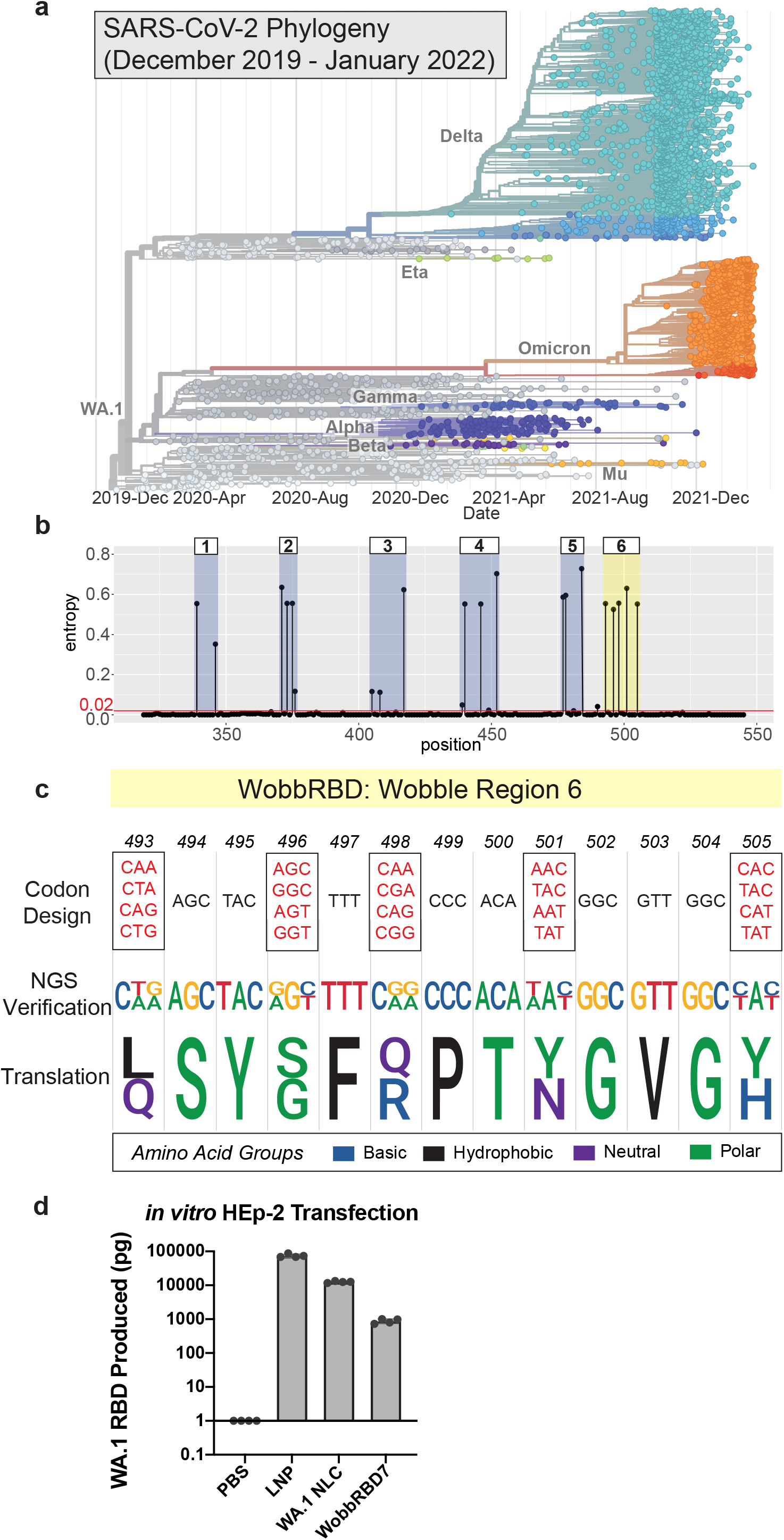
WobbRBD library design. A) Phylogenetic tree of SARS-CoV-2 spike protein sequence diversity from December 2019 to January 2022 developed using the Nextstrain ncov GISAID global dataset. B) Entropy plot of the normalized Shannon entropy at each amino acid position in the SARS-CoV-2 spike RBD using Nextstrain data from the timeline shown in (A). Entropy cut-off is set at 0.02. Wobble regions are highlighted and numbered 1-6. C) An example of codon design, DNA library verification, and anticipated protein library sequence for wobble region 6. Codons are designed to encapsulate strain diversity using a limited number of nucleotide changes in library creation. Library sequences were verified by next-generation sequencing and are shown as the relative probabilities of each base at each position. When translated, these 1000+ sequences match the predicted amino acid diversity. D) Hep2 cells were transfected with 1 µg non-amplifying mRNA-LNP encoding secreted WA.1 RBD, 1 µg saRNA-NLC encoding secreted WA.1 RBD, 1 µg saRNA-NLC encoding secreted RBDs with all wobble sites (WobbRBD7), or a PBS control. Supernatants were harvested after overnight incubation at 37ºC, 3 replicate wells were pooled and concentrated, and RBD production was determined by ELISA. Any readings below 1 pg were set to this value to enable visualization. Points shown are technical replicates from two dilutions run in duplicate.

Flex DNA libraries were cloned into a self-amplifying vector utilizing Venezuelan Equine Encephalitis Virus (VEEV) replication machinery as previously published^15^. Plasmid libraries were then transcribed into self-amplifying mRNA (saRNA) and complexed with nanostructured lipid carriers (NLCs), a thermostable *in vivo* RNA delivery system alternative to the traditional lipid nanoparticle (LNP) with demonstrated high efficacy in preclinical studies^15-18^. NLCs comprise a highly thermostable emulsion consisting of a solid lipid (dynasan) and liquid oil (squalene) core and the cationic lipid 1,2-dioleoyl-3-trimethylammonium propane (DOTAP), emulsified with Tween and Span surfactants and to which RNA binds on the exterior of the particle where it is stabilized and effectively delivered to cells with relative protection from RNase challenge (summarized in Table S2)^15, 19^. Successful and functional formulation was additionally verified through *in vitro* transfection of HEp-2 cells and detection of secreted WA.1 RBD antigen (Fig. 2D). Importantly, WobbRBD7 encodes diverse RBD sequences that diverge from WA.1 and may not be fully detected by the utilized ELISA capture antibodies.

The WobbRBD library encompasses the initial emergence of SARS-CoV-2 until the emergence of the first Omicron variant in January 2022 (Fig 2A). To assess the predictive power of this entropy-based approach, individual flex sites were assessed for diversity in ‘future’ strains. Taking into account the full 5+ year mutational history of the SARS-CoV-2 RBD, wobble regions designed for the 2022-based WobbRBD library still include around two-thirds of the 2025 mutational load (29/44 sites; Fig. S1A). At the time of design, the RBD entropic cut-off of 0.02 designated 23 sites of high entropy of which 16 were noted mutations in VOC. Of the remaining seven sites, five (positions 346, 376, 405, 408, and 481) became increasingly prevalent in the ensuing Omicron family of variants, including BA.5, XBB.1.5, JN.1, and XFG, based on the mutation-tracking data available through CoV-Spectrum^20^ (Fig. S1B).

### Early Innate Response Comparison Across Vaccine Modalities

To determine the immunogenicity of wobble vaccines in comparison to traditional modalities, a kinetic analysis of vaccine recipients was performed using high-dimensional spectral flow cytometry to identify 26 distinct immune subsets within the draining LNs. The WobbRBD7 saRNA-NLC library was tested alongside protein-Alum, non-replicating mRNA-LNP vaccines, and a directly comparable single strain saRNA-NLC control. All utilized or encoded the WA.1 RBD. All four modalities were compared across a two-week response time course in mice following intramuscular (i.m.) vaccination in the calf through analysis of the popliteal and inguinal draining lymph nodes (dLNs) by flow cytometry (gating shown in Fig. S2 and S3). No significant weight decrease or other adverse effects were observed due to treatment (Fig. S4A).

At three days post-vaccination, a mounting immune response can be detected through a shift in the immune composition of the dLNs. Using immune subpopulations as features, mRNA-LNP, WA.1 saRNA-NLC, and WobbRBD7 saRNA-NLC vaccinated mice can be clearly separated from PBS control and Alum-vaccinated animals via a principal component analysis (PCA) of the innate and adaptive responses (Fig. 3A). Innate recruitment and activation play a pivotal role in early immune activation within the dLN; notably, there is a significant increase in the frequency of the overall innate compartment in the WobbRBD7 group compared to the PBS control (Fig. S4B; *F*_4, 20_ = 4.597, *p* = 0.0085). Key innate and adaptive immune populations characterized the three most potent vaccine modalities compared to the PBS and Alum conditions, with hierarchical clustering establishing groupings of features (Fig. 3B).

**Figure 3.**
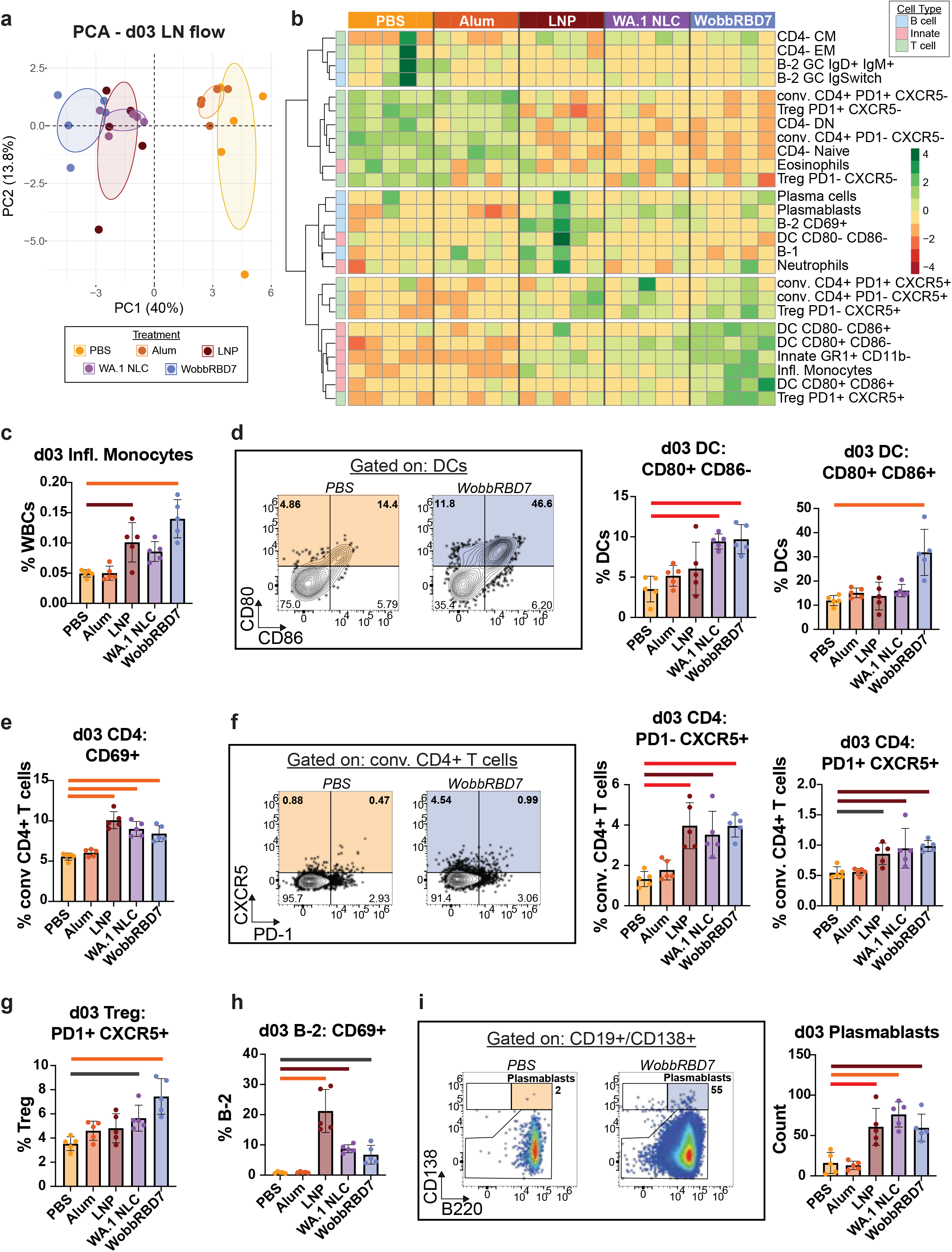
Comparison of early immune responses by vaccine modality. C57BL/6 mice were vaccinated i.m. with 20 μl containing either PBS (yellow), 5 μg WA.1 RBD protein in a 1:1 Alum solution (orange), 1 μg WA.1 RBD mRNA-LNP (brown), 1 μg WA.1 RBD saRNA-NLC (purple), or 1 μg WobbRBD7 saRNA-NLC (blue). The popliteal and inguinal draining lymph nodes were collected and analyzed by flow cytometry at three days post-vaccination. Pre-gating is shown in Fig. S2. Significance was determined by one-way ANOVA through comparison to the PBS control group with correction for multiple comparisons. A) PCA plot of key populations identified by flow cytometry at day 3. All populations were expressed as a percent of white blood cells (WBCs). B) Heatmap of the populations used to create (A), compared between vaccination conditions and individual samples (by column). Data is scaled by row and clustered using the Ward.D2 method from the R ‘pheatmap’ library. (C-I) Flow plots show the relevant gates for the PBS (yellow) and WobbRBD7 saRNA-NLC (blue) with the relevant percent of the parent population. n = 5 mice per group. No bar = n.s., black bar (*) = *p* ≤0.05; crimson bar (**) = *p* ≤0.01; red bar (**) = *p* ≤0.001; orange bar (****) = *p* ≤0.0001. Bar plots display the mean with error bars for standard deviation.

Within the innate compartment, unique shifts were identified in the inflammatory monocyte, dendritic cell (DC), and a potential lymphoid tissue inducer (LTi) or innate lymphoid cell (ILC) populations. The mRNA-LNP and WobbRBD7 saRNA-NLC conditions show a significant increase in inflammatory monocytes (GR1+ CD11b+) relative to the PBS controls and were verified to express Ly6C rather than Ly6G (Fig. 3C; *F*_4, 20_ = 14.41, *p* < 0.0001). Additionally, the LTi or ILC population lacks distinctive cell type markers (TCRb-CD19-CD11b-CD161-), expresses several Th1-like response markers (IL7R+ CD62L+ CXCR3+ CD44+ Ly6C+), and shows a significant increase in the mRNA-LNP and saRNA-NLC conditions (Fig. S4C; *F*_4, 20_ = 22.60, *p* < 0.0001). Unique to the saRNA-NLC conditions, DCs additionally bolster these immune responses through an increased costimulatory phenotype. Intriguingly, CD80+ CD86-DCs are more abundant in both the WA.1 and WobbRBD7 saRNA-NLC conditions (*F*_4, 20_ = 9.579, *p* = 0.0002), whereas CD80+ CD86+ double-positive DCs are specifically increased in the WobbRBD7 group alone (*F*_4, 20_ = 11.66, *p* < 0.0001), suggesting a potential difference in the specific expression of these similar costimulatory molecules (Fig. 3D). Together, these data show qualitative, as well as quantitative, shifts in the innate response to both saRNA-NLC-delivered vaccines compared to other vaccine modalities.

### Early Adaptive Response Comparison Across Vaccine Modalities

Under homeostatic conditions, LNs contain ∼80% T cells; however, vaccination induces short-term retention and a significant expansion of the B cell compartment in the dLNs, which was detected under the mRNA-LNP, WA.1 saRNA-NLC, and WobbRBD7 saRNA-NLC conditions to produce closer to a 50:50 B:T cell ratio (Fig. S4D; *F*_4, 20_ = 17.59, *p* < 0.0001). Under further inspection, additional adaptive immune features established distinctive characterization of different vaccine modalities here. Activated CD4+ T cell responses distinguished the saRNA-NLC conditions, while an early B cell-driven response with prominent CD69 expression characterized the non-amplifying mRNA-LNP-induced response (Fig. 3B).

In the CD4+ T cell compartment, mRNA-LNP and saRNA-NLC conditions induced CD4+ T cell activation and altered their localization within the dLN. Across the mRNA-LNP and saRNA-NLC conditions, this was shown through increased CD69 expression for LN retention (Fig. 3E; *F*_4, 20_ = 28.34, *p* < 0.0001), as well as upregulation of CXCR5 to promote migration to the T-B border (Fig. 3F). This increase in follicular localization was detected in both recently activated CD4+ T cells (CXCR5+ PD1-; *F*_4, 20_ = 11.99, *p* < 0.0001) and ones with a more typical follicular helper T cell (Tfh) phenotype (CXCR5+ PD1+; *F*_4, 20_ = 7.092, *p* = 0.0010) across the mRNA-LNP and saRNA-NLC conditions (Fig. 3F). Specific to the saRNA-NLC conditions, within the CD25+ IL7R-regulatory T cell (Treg) population, CXCR5+ PD1+ follicular regulatory T cells (Tfr) were more abundant (Fig. 3G; *F*_4, 20_ = 9.306, *p* = 0.0002).

Additionally, in the WobbRBD7 saRNA-NLC condition alone, there was 1) an overall decrease in CD62L MFI in the conventional CD4+ T cell compartment (Fig. S4E; *F*_4, 20_ = 9.246, *p* = 0.0002) and 2) the emergence of a distinct CD4+ T cell population expressing a modified CD45R that bound the anti-B220 antibody clone RA3-6B2 (Fig. S4F; *F*_4, 20_ = 13.68, *p* < 0.0001). This behavior has been previously observed in murine CD4+ T cells at a similar timeframe post-activation from certain stimuli^21^, which suggests a possible shift in glycosylation from a naïve CD45R state that is not bound by this antibody. In other models, B220+ CD4+ T cells have been reported to be linked to a post-proliferation, pre-apoptotic state^22, 23^; however, their exact role here is unknown, and this population was not observed at later time points.

Similar to the overall activation observed across CD4+ T cells, signs of activation and retention were observed within the expanded B cell compartment. Across the mRNA-LNP and saRNA-NLC conditions, an increased percentage of B cells expressed CD69, with the most extreme phenotype in the non-amplifying mRNA-LNP condition (Fig. 3H; *F*_4, 20_ = 27.82, *p* < 0.0001). This prominent feature of the mRNA-LNP condition might be explained by the 10-fold difference in short-term antigen production observed in Fig. 2D between the mRNA-LNP and the saRNA-WA.1 NLC conditions. No meaningful detection of germinal center (GC) B cells was observed at this early timepoint despite the increase in Tfh and Tfr within the dLNs; however, plasmablast accumulation was identified, likely derived through early extrafollicular-driven differentiation as has been reported previously^24^ (Fig. 3I; *F*_4, 20_ = 16.48, *p* < 0.0001). Specifically, within the WobbRBD7 saRNA-NLC condition, as above, B cells demonstrate an overall decrease in CD62L MFI (Fig. S4G; *F*_4, 20_ = 23.36, *p* < 0.0001). Overall, these facets describe and distinguish between mRNA-LNP and saRNA-NLC-stimulated early immune responses and demonstrate consistency between the single strain and wobbled saRNA-NLC conditions with a few unique features of the wobble vaccine modality.

### Peak Adaptive Response Comparison Across Vaccine Modalities

Next, we investigated how the above differences in early vaccine responses alter the peak adaptive immune response at two weeks post-vaccination. PCA analysis of immune composition revealed ongoing intensity in saRNA-NLC-induced immune response compared to the PBS, Alum, and mRNA-LNP responses, which have waned by d14 (Fig. 4A), driven by a distinct immune cassette of activated T and B cells (Fig. 4B).

**Figure 4.**
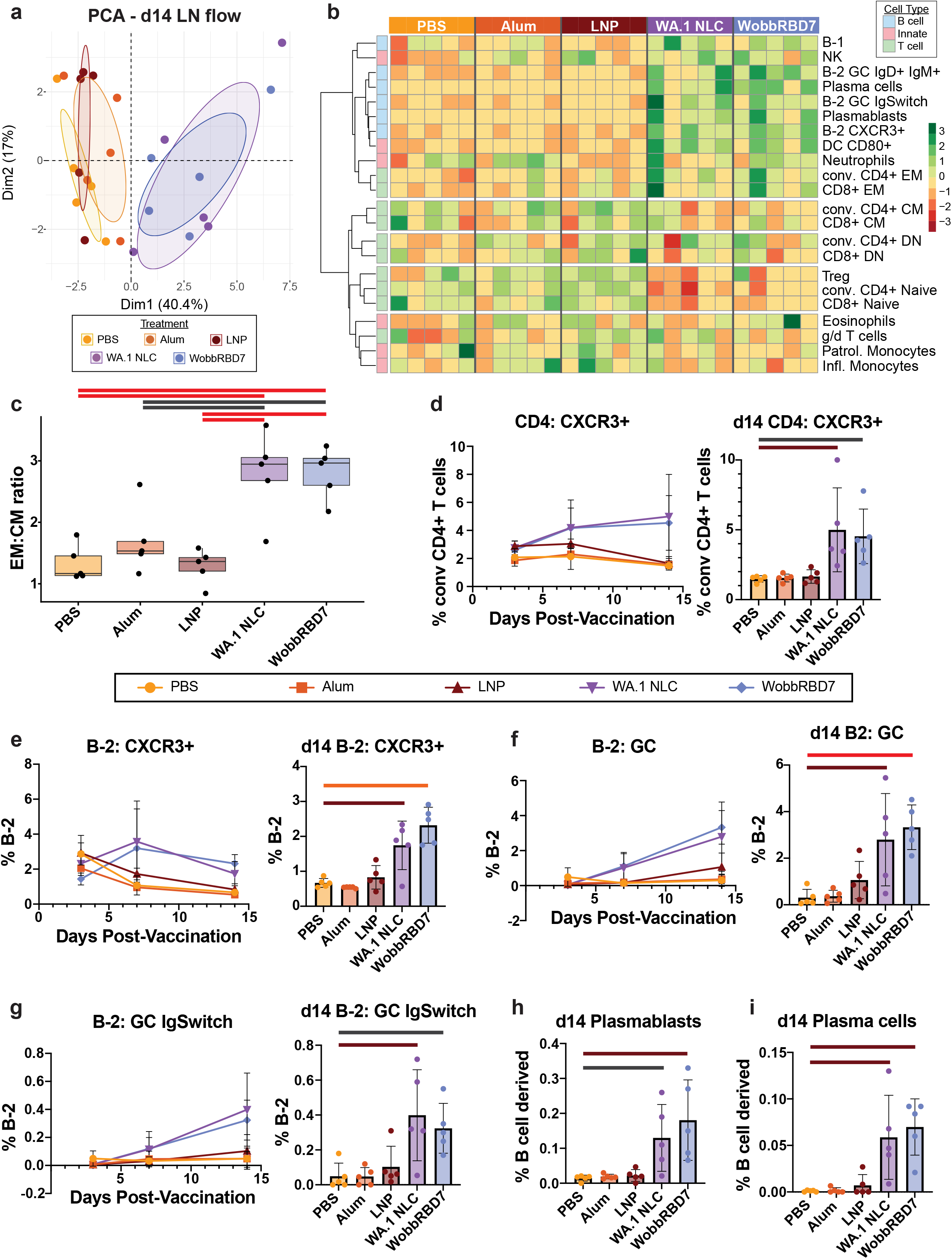
Comparison of peak adaptive vaccine responses by vaccine modality. C57BL/6 mice were vaccinated i.m. with 20 μl containing either PBS (yellow), 2 μg WA.1 RBD protein in a 1:1 Alum solution (orange), 1 μg WA.1 RBD mRNA-LNP (brown), 1 μg WA.1 RBD saRNA-NLC (purple), or 1 μg WobbRBD7 saRNA-NLC (blue). Popliteal and inguinal draining lymph nodes were collected and analyzed by flow cytometry at days 7 and 14 post-vaccination in comparison to the day 3 data shown in Fig. 3. The gating scheme for days 7 and 14 data is shown in Fig. S3. Significance was determined by one-way ANOVA through comparison to the PBS control group with correction for multiple comparisons. A) PCA plot of key populations identified by flow cytometry at day 14. All populations were expressed as a percent of WBCs. B) Heatmap of the populations used to create (A), compared between vaccination conditions and individual samples (by column). Data is scaled by row and clustered using the Ward.D2 method from the R ‘pheatmap’ library. C) Boxplot of the ratio of conventional CD4+ T cell effector memory (EM; CD44+ CD62L-) and central memory (CM; CD44+ CD62L+) populations at day 14 in the dLNs by vaccination condition. All ratios are calculated from percent of WBC of the relevant population. (D-G) Data is shown both as a time course for days 3, 7, and 14, and with the day 14 data depicted as a separate bar graph. (H-I) Results at day 14 are shown as bar graphs as described above. n = 5 mice per group for days 3 and 14, n = 3 mice per group for day 7. No bar = n.s., black bar (*) = *p* ≤0.05; crimson bar (**) = *p* ≤0.01; red bar (**) = *p* ≤0.001; orange bar (****) = *p* ≤0.0001. Bar plots and the points on time course plots display the mean with error bars for standard deviation.

A significant increase in the ratio of effector memory (EM) to central memory (CM) CD4+ T cells distinguishes the saRNA-NLC conditions from other vaccination modalities and denotes increased blood-based localization of activated T cells (Fig. 4C; *F*_4, 20_ = 12.85, *p* < 0.0001). CD4+ T cells (*F*_4, 20_ = 5.890, *p* = 0.0027) and B cells (*F*_4, 20_ = 17.08, *p* < 0.0001) also both show increased expression of CXCR3 under the WA.1 saRNA-NLC and WobbRBD7 saRNA-NLC conditions, indicating activation of an IFN-γ-driven microenvironment (Fig. 4D-E).

These T-B interactions and increased exposure to antigen contribute to the establishment of GCs within the B cell follicles, which are crucial for affinity maturation in vaccination responses^25, 26^. The WA.1 and WobbRBD7 saRNA-NLC conditions both show a significant increase in GC B cells at day 14 (Fig. 4F; *F*_4, 20_ = 8.698, *p* = 0.0003), with an additional increase in the frequency of class-switched GC B cells (Fig. 4G; *F*_4, 20_ = 6.158, *p* = 0.0021). Correspondingly, there is a significant increase in plasmablast (Fig. 4H; *F*_4, 20_ = 6.482, *p* = 0.0016) and plasma cell (Fig. 4I; *F*_4, 20_ = 9.182, *p* = 0.0002) frequencies within the dLNs under the saRNA-NLC conditions specifically, denoting the potency of this vaccination platform. These similarities in potent B cell stimulation by day 14 assuage concerns about the impact of introduced heterogeneity in the WobbRBD7 library and demonstrate the overall efficacy of the saRNA-NLC vaccine format.

### Enhanced Production of Broadly-Binding Antibodies

With the confidence that wobble vaccines do not impair overall vaccine responses and may enhance or accelerate activation at certain stages, we next examined whether WobbRBD can effectively alter epitope targeting of the humoral response. The overall magnitude and specificity of the resulting antibody responses were assessed through a panel of SARS-CoV-2 variant spikes (MesoScale Discovery). Both the WA.1 and WobbRBD7 saRNA-NLC conditions produced a significant increase in IgG titer against the WA.1 spike by day 14, more potent than any of the above vaccination modalities (Fig. 5A; *F*_4, 20_ = 54.35, *p* < 0.0001).

**Figure 5.**
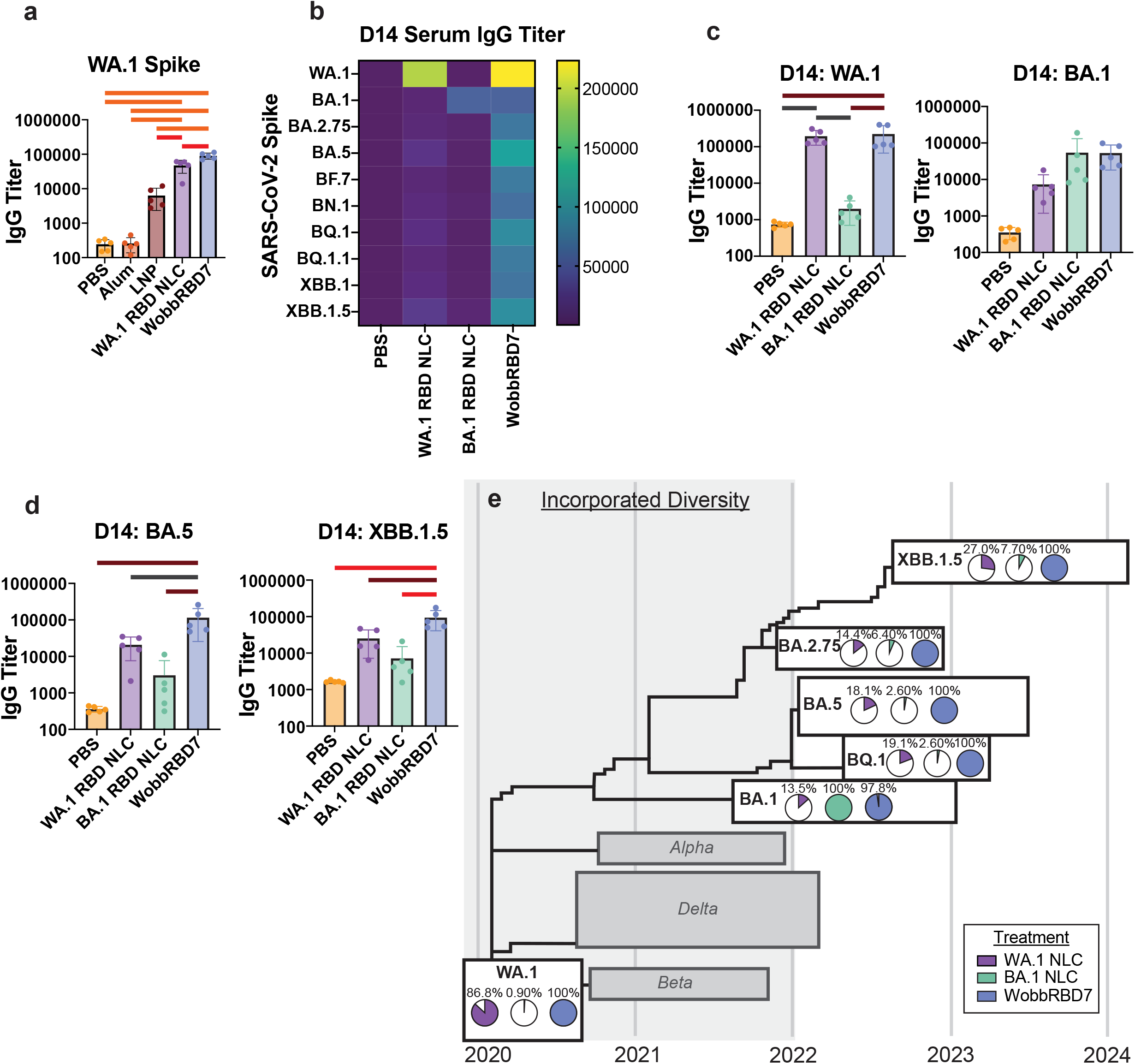
WobbRBD promotes increased antibody breadth. A) C57BL/6 mice were vaccinated i.m. with 20 µl containing either PBS (yellow), 2 µg WA.1 RBD protein in a 1:1 Alum solution (orange), 1 µg WA.1 RBD mRNA-LNP (brown), 1 µg WA.1 RBD saRNA-NLC (purple), or 1 µg WobbRBD7 saRNA-NLC (blue). Serum was collected at day 14 post-vaccination and evaluated for SARS-CoV-2 spike binding using the MSD V-PLEX SARS-CoV-2 Panel 34 of variant spike proteins. IgG titers are shown in relative units. B-E) C57BL/6 mice were vaccinated i.m. with 20 µl containing either PBS (yellow), 1 µg WA.1 RBD saRNA-NLC (purple), 1 µg Omicron BA.1 RBD saRNA-NLC (teal), or 1 µg WobbRBD7 saRNA-NLC (blue). Serum was collected and evaluated as in (A). Spike binding data presented as a heatmap. C-D) Spike binding data presented as bar plots. E) Spike binding data presented as percent of maximum mean IgG titer by variant, laid out along a sketch of a phylogeny timeline. The original phylogeny was generated using Nextstrain. n = 5 mice per group. No bar = n.s., black bar (*) = p ≤ 0.05; crimson bar (**) = p ≤ 0.01; red bar (**) = p ≤ 0.001; orange bar (****) = p ≤ 0.0001. Bars display the mean with error bars for standard deviation.

Having documented that the saRNA-NLC format induced potent immune responses, the following experiment compared WobbRBD7 to two single strain saRNA-NLC vaccines at either end of its initial design timeline: WA.1 (2019) and Omicron BA.1 (early 2022). Both the WA.1 and BA.1 RBD saRNA-NLCs induced qualitative increases in anti-spike IgG antibodies against their respective single strain spike, with limited cross-reactivity to the panel of additional variant spikes (Fig. 5B-C). This potency is reflected in significant differences in WA.1 RBD titer (*F*_3, 16_ = 9.156, *p* = 0.0009) induced by treatment but no significant changes in BA.1 RBD titer across all treatments, perhaps due to variability in serum titers (*F*_3, 16_ = 2.330, *p* = 0.1130). Nonetheless, similar patterns are observed in the shifts in BA.1 RBD binding compared to WA.1. WobbRBD7 saRNA-NLC, on the other hand, produced both comparable antibody binding responses to the WA.1 and BA.1 saRNA-NLCs against their respective spikes and was the only condition to maintain similar binding efficacy across the panel of variant spikes (Fig. 5B-D; Fig. S5), including BA.5 (*F*_3, 16_ = 7.175, *p* = 0.0029) and XBB.1.5 (*F*_3, 16_ = 11.35, *p* = 0.0003).

This increased breadth is made more impactful when we consider the timeline of SARS-CoV-2 spike mutations on display. Many of these variants (e.g. BA.2.75, BA.5, BQ.1, XBB.1.5) emerged after the timeline incorporated in the construction of the WobbRBD7 library (Fig. S1B) and yet WobbRBD7 saRNA-NLC produced the maximum IgG-binding signal of the three saRNA-NLC conditions to all post-January 2022 variants tested here (Fig. 5E). Altogether, these data suggest that while wobble vaccine design elicits similar immunogenicity to single-strain controls, the use of RES in library design drives substantially increased breadth in cross-strain targeting.

## Discussion

Wobble vaccines represent a promising and innovative investigation into cocktail vaccine design with demonstrated efficacy in increasing variant-binding antibody breadth. The entropic analysis applied here has demonstrated its utility in the prediction of emerging regions of heterogeneity (when informed by sufficient depth of sequencing and pathogen monitoring). We have shown that the introduction of these decoupled mutations to produce a heterogeneous antigen pool does not disrupt a typical vaccination immune response and may serve to augment it in ways that require further investigation. Similarly, IgG titers against ancestral antigen were preserved, and increased binding against a wider breadth of variants suggested an increase in the targeting of conserved epitopes. This work builds upon the demonstrated efficacy of multi-strain hemagglutinin influenza studies^10, 27^ and helps to elucidate some of the underlying features of RES that may govern both sets of work, in addition to far-reaching but similar phenomena, such as HLA-targeting in transplants^28^. Wobble vaccines seek to innovate upon this efficacy using a robust mRNA-based platform to allow for 1) streamlined and cost-effective production of diverse antigen, and 2) generation of a potent immune response in a single immunization. These factors are key for large-scale distribution of such a vaccine and ease of administration, which combined with the increased breadth of protection, can serve to increase the effective lifetime of vaccines and counteract the time between a vaccine’s inception, development, and deployment.

Additionally, on its own, the vaccine modality comparison offers an in-depth look at novel characteristics of immune activation within a draining LN. In general, saRNA-NLC vaccines were more potent in inducing sustained immune activation at two-weeks post-vaccination through an increase in germinal center B cells, ASCs, and CXCR3 expression across CD4+ T cells and B cells (Fig. 4). Interestingly, the wobble construct drove specific differences shortly after vaccine administration through enhanced costimulatory molecule expression on DCs, decreased CD62L expression on CD4+ T cells and B cells, and B220-like modifications to CD45R on CD4+ T cells, which had waned after two weeks (Figs. 3 & S4). These early differences could potentially be related to variable concentrations of mutated antigens leading to populations of lymphocytes activated by either high or low levels of antigen with distinct resulting phenotypes. Additionally, later timepoints may point to the presence of key microenvironments for B and T cell organization and crosstalk, as CXCR3 is used both for localization to sites of inflammation from the blood as well as specialized localization within the LN. It is known that CXCR3 is vital for DC-CD4+ T cell interactions within the dLN^29, 30^ and could possibly point to B cell localization to these areas as well. Currently the exact cause of these changes is not clear and will be important to answer in subsequent studies through more direct investigation of the emerging LN microenvironments and the effects of variable levels of antigen stimulation *in vivo*.

While the current work demonstrates the potential of this wobble vaccine approach against SARS-CoV-2, wobble vaccines stand to benefit multiple vaccination efforts across several goals. Seasonal pathogens, such as influenza A and B, already have large databases of global historic sequencing information to shape a similar vaccine. Highly mutable pathogens, such as human immunodeficiency virus (HIV), represent another clear target, with the potential for either personalized medical approaches to craft vaccines catered to an individual’s mutational load or widely-applicable preventative vaccines focused on founder strains that establish initial infections. With sufficient *in silico* or animal model testing for escape mutations, wobble vaccines even have potential for the quick and effective creation of proactive vaccines against novel emerging pathogens.

Further experimentation remains to fully evaluate the potential and long-term protection capabilities of wobble vaccines, either in saRNA-NLC or other mRNA-based vaccine modalities. One question of high concern is whether an approach like RES could manage to circumvent antigenic imprinting where the first memory developed against a pathogen shapes what can be responded to in future variant encounters and may make potent variant-specific responses more difficult to mount. Another question might center around how these efforts to shape epitope targeting can enhance or interfere with the development of neutralizing antibodies. Investigation of how much diversity can be encoded in this vaccine approach is key to investigations toward highly mutable pathogens like influenza. Finally, the effect of cocktail vaccines like these on memory formation and recall events invites additional study^31^. With a wealth of potential applications and avenues for further exploration, this initial demonstration illustrates the potential for wobble vaccines as a method to shape antibody-epitope responses in a proactive manner to confer long-lived protection.

## Methods

### Library Design

The WobbRBD1-7 libraries were constructed based on normalized Shannon entropy data on the SARS-CoV-2 RBD taken from December 2019 through January 2022. Entropy data, as well as all phylogenetic trees shown here, were derived from the Nextstrain ncov GISAID global dataset. R and ggplot were used to visualize the entropy distribution. A cutoff of 0.02 to designate ‘high entropy’ amino acid sites was designated to encapsulate a reasonable amount of the demonstrated heterogeneity while following guidelines for the construction of the DNA library. Appropriate nucleic acid base substitutions to allow for the minimum number of possible bases while encoding all possible amino acid variations were determined by hand. WA.1 and BA.1 RBD sequences were acquired from GenBank. An IgK secretion signal was added upstream of the single strain and wobbled RBD sequence constructs to induce secretion of the produced protein. DNA libraries were produced by GenScript and verified by next-generation sequencing. Sequences are listed in Table S1.

### saRNA-NLC Formulation

#### *In vitro* transcribed (IVT) saRNA synthesis

DNA libraries were cloned into VEEV-based saRNA backbone plasmids using standard cloning techniques. *In vitro* transcription of saRNA from DNA template libraries was conducted as previously described^15^. Briefly, DNA templates for saRNA production were linearized with NotI-HF (New England Biolabs), purified by phenol-chloroform extraction, and vaccine saRNA produced by IVT using T7 polymerase, RNase inhibitor and pyrophosphatase (Aldevron). DNA was digested away (DNase I, Aldevron) and Cap0 structures added using guanylyltranferase (Aldevron). A poly-A tail sequence was encoded by the saRNA template removing any need for enzymatic tailing.

Produced saRNA was purified by lithium chloride precipitation, washing, and resuspension prior to aliquoting and storage at -80°C. Agarose gel electrophoresis was used to characterize saRNA integrity, and RNA concentration assessed by UV absorbance (NanoDrop 1000) and RiboGreen assay (ThermoFisher).

#### saRNA-NLC production

Nanostructured lipid carriers were manufactured in bulk as previously described^15, 18^. Briefly, squalene, DOTAP (N-[1-(2,3-dioleoyloxy)propyl]-N,N,N-trimethylammonium chloride), sorbitan monostearate (Span 60) and glyceryl trimyristate (Dynasan 114) were mixed and heated in a bath sonicator to create an oil phase, which was then emulsified with a 10mM sodium citrate trihydrate solution containing polysorbate 80 (Tween 80) using a high-speed emulsifier (Silverson Machines) to create a crude emulsion that was subsequently homogenized at 30,000 psi in a laboratory microfluidizer (Microfluidics M110P) for approximately 10 discrete passes until the desired particle size was achieved. The NLC product was finally filtered through a 0.22µm polyethersulfone filter and stored aliquoted at 2-8°C in single-use glass vials until complexing with mRNA.

### mRNA-LNP Formulation

#### *In vitro* transcribed mRNA synthesis

The SARS-CoV-2 WA.1 RBD single strain construct from Table S1 was synthesized as mRNA for the cargo of standard mRNA-LNPs, as previously described^32, 33^. For *in vitro* transcription, plasmids were linearized with Not-I HF (New England Biolabs) overnight at 37 °C. Linearized templates were purified by sodium acetate (Thermo Fisher Scientific) precipitation and rehydrated with nuclease-free water. IVT was performed at 37 °C using the HiScribe T7 RNA Polymerase (Aldevron) following the manufacturer’s instructions (N1-methyl-pseudouridine modified). The resulting RNA was treated with DNase I (Aldevron) for 30 min to remove the template and was then purified using lithium chloride precipitation (Thermo Fisher Scientific). The RNA was heat denatured at 65 °C for 10 min before capping with a type 1 cap structure using guanylyl transferase and 2′-O-methyltransferase (Aldevron). mRNA was then enzymatically tailed with poly-A polymerase (Aldevron) and phosphatase (NEB) treated before a final purification. mRNA concentration was measured using a Nanodrop. Purified mRNA products were analyzed by capillary gel electrophoresis (Agilent Fragment Analyzer) to ensure purity.

#### mRNA-LNP production

The mRNAs were diluted in 10 mM citrate buffer (pH 3) to create the aqueous phases. To prepare the organic phase, SM-102 (Broadpharm), cholesterol (Sigma), DMG-PEG-2k (Avanti Polar Lipids), and DSPC (Avanti Polar Lipids) were added to 100% ethanol at a ratio of 50:38.5:1.5:10. A mass ratio of 20:1 (lipid/mRNA) was used for all formulations. The two phases were mixed using a NanoAssemblr benchtop device containing a microfluidic cartridge (Precision NanoSystems Inc.) at an aqueous to organic flow rate ratio of 3:1 at 12 mL/min. The LNPs were next diluted 6× in 10 mM Tris buffer and concentrated using 30 kDa MWCO centrifugal filters (Millipore Sigma), followed by a 7× diafiltration with 6% (w/v) sucrose. After sterile filtering, particle size was determined by dynamic light scattering on a Zetasizer Pro (Red) (Malvern Panalytical). A RiboGreen assay (Invitrogen) was performed to calculate the encapsulation percentage and concentration of mRNA cargos. After size and concentration were characterized, LNP solutions were diluted with 6% sucrose to the desired final concentration, aliquoted, and stored at −80 °C until use.

### In vitro *Transfection*

HEp-2 cells were cultured in Eagle’s Minimum Essential Media with 10% FBS and incubated in a VWR incubator at 37 °C and 5% CO2. Cells were passaged 1:10 every 3 days for maintenance. For transfection, cells were split 1:2 from confluency at 1 ml/well into a 24 well plate. Wells were transfected with 1 μg of either the above mRNA-LNP, WA.1 saRNA-NLC, or WobbRBD7 saRNA-NLC constructs, or a PBS control in technical triplicate. After overnight incubation, the supernatant was removed and any remaining cells were pelleted at 350 xg for 10 min. The collected supernatant was pooled within a condition and underwent concentration and buffer exchange to PBS in a 10 kDa concentrator. RBD production was quantified using an RBD ELISA kit (Invitrogen) according to manufacturer protocol (see Key Resources Table).

### Mice

C57BL/6 female mice were received at 6 weeks of age from the Jackson Laboratory. All mice used in the study were female for purposes of reproducibility and were used for experimentation between 6 and 10 weeks of age.

### Vaccination

Mice were vaccinated intramuscularly in the calf using 0.3 ml insulin syringes containing 20 μl doses. mRNA-LNP and saRNA-NLC formulations were used at 1 μg per mouse. Protein-Alum doses were either 2 μg or 5 μg of WA.1 RBD in a 1:1 mixture of PBS:Alhydrogel (see Key Resources Table).

### Serum Collection and Processing

Blood samples were collected at two weeks post-vaccination through submandibular venipuncture (or cheek bleeds) using a 5 mm lancet to collect 50-100 μl. Serum was isolated through 30 minute incubation at room temperature followed by centrifugation at 3000 xg for 10 minutes. Supernatant was stored at either 4 °C or -80 °C for short-or long-term storage respectively.

### Flow Cytometry

For lymph node collection, mice were euthanized by carbon dioxide asphyxiation followed by cervical dislocation. Popliteal and inguinal lymph nodes were collected at either 3, 7, or 14 days post-vaccination into 200 μl PBS and homogenized using pestles in 1.5 ml Eppendorf tubes. After allowing large chunks to settle, the supernatant was transferred to a Costar V-bottom 96-well plate and stained with viability dye in PBS, washed with FACS buffer (PBS with 2% FBS and 1 μM EDTA), and stained with one of the two following panels of antibodies and viability dye listed in the Key Resources table. After staining, samples were washed and filtered through a 35 μm filter into 5 ml FACS tubes. Samples were analyzed on a 5-laser Cytek Aurora using Cytek SpectroFlo software in the Emory Pediatrics/Winship Flow Cytometry Core (ECFCC). Further analysis was run using FlowJo v10.10.0.

### Serum Antibody Binding

Quantification of serum antibodies was performed using multiplex electrochemiluminescence serology assays by Meso Scale Discovery (MSD), as previously described^34-36^. Serum spike-specific IgG antibodies were measured using V-Plex SARS-CoV-2 Panel 34 (kindly provided by MSD), according to the manufacturer’s instructions. Briefly, antigen-specific plates were blocked using MSD blocker at room temperature for 30 minutes with shaking at 700 rpm. The samples were then diluted 1:500, 1:5,000, and 1:50,000, incubated on the plates for 2 hours at room temperature, and detected by SULFO-TAG–conjugated goat anti-mouse IgG antibody (Meso Scale Discovery). The plates were subsequently washed with 1X MSD wash buffer, followed by the addition of MSD Gold Read Buffer B to each well. Plates were washed 3 times with MSD wash buffer after each stage. Optical densities were measured using an MSD plate reader, and the results were analyzed with Discovery Workbench software (version 4.0) with antibody levels reported in arbitrary units per mL against SARS-CoV-2 spikes.

### Statistics

All statistical comparisons were run in Graphpad Prism (v10.6.0) using one-way ANOVA tests with multiple comparisons. Comparisons with *p* values ≤0.05 were considered significant and reported based on the color bars shown in figures and described in the figure captions: no bar = n.s., black bar (*) = *p* ≤0.05; crimson bar (**) = *p* ≤0.01; red bar (**) = *p* ≤0.001; orange bar (****) = *p* ≤0.0001.

### Software and Analysis

R (v4.4.1, release 14 June 2024) and Graphpad Prism (v10.6.0) were used for computational analysis. Heatmaps were generated using the ‘pheatmap’ library (v1.0.13) using relevant non-redundant flow cytometry populations as a percent of WBC. Clustering was performed using Ward’s method (Ward.D2). Entropy analyses were run using the ‘tidyverse’ library and visualized using ‘ggplot2.’ Sequence alignments and translation were carried out using the ‘msa’ library and visualized using the ‘ggseqlogo’ library. PCA analysis and plots were generated using the ‘factoextra’ library. Statistical analysis was run in Prism. Adobe Illustrator was used for further figure creation and post-processing.

## Acknowledgements

This work was supported by the Emory Pediatrics/Winship Flow Cytometry Core (ECFCC) for the facilities and instruments used to run all flow cytometry data presented here. This work was also supported by the production of WA.1 RBD protein by the Frederick National Laboratory as part of SeroNet. We acknowledge funding from the DOD (W81XWH-22-1-0572) for supporting this work.

## Author Contributions

**Peter R. McIlroy**: investigation (lead), formal analysis (lead), visualization (lead), writing – original draft preparation (lead), writing - review and editing (equal). **Matthew C. Woodruff**: conceptualization (lead), funding acquisition (lead), methodology (lead), project administration (lead), supervision (lead), investigation (supporting), visualization (supporting), writing – review and editing (equal). **Wendy M. Zinzow-Kramer**: methodology (supporting), investigation (supporting), project administration, writing – review and editing (supporting). **Madison L. Ellis:** investigation (supporting), formal analysis (supporting). **Mehul S. Suthar**: formal analysis (supporting), supervision, writing – review and editing (supporting). **Eduard Melief**: conceptualization (supporting), resources (equal), validation (equal). **Emily A. Voigt**: conceptualization (supporting), resources (equal), validation (equal), writing – review and editing (supporting). **Hannah E. Peck**: resources (equal), validation (equal), writing – review and editing (supporting). **Loren E. Sasser**: resources (equal), validation (equal). **Daryll Vanover**: resources (equal), validation (equal), methodology (supporting). **Philip J. Santangelo**: resources (equal), validation (equal), supervision (supporting). **Mohammad Ali**: methodology (supporting), writing – review and editing (supporting).

## Data Availability

Raw data are available upon reasonable request to the authors.

## Materials & Correspondence

Correspondence or requests for materials should be addressed to Matthew C. Woodruff.

