## Supplementary Figures and Tables for "Wobble Vaccines: Complex Vaccine Antigen Pools Promote Increased Antibody Breadth and Cross-Strain Viral Targeting in SARS-CoV-2"

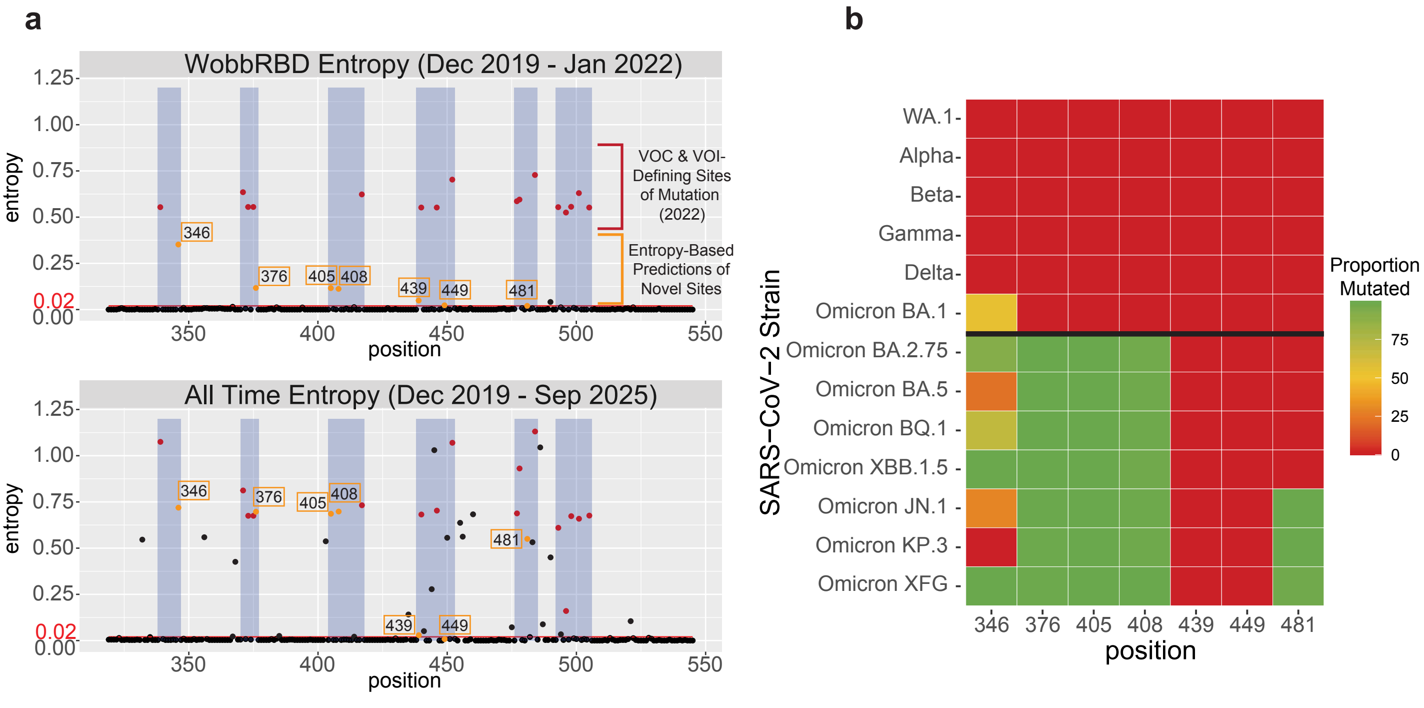

**Supplementary Figure 1. Analysis of captured diversity by WobbRBD entropy model –**
A) Plots of the Shannon normalized entropy for the WobbRBD using SARS-CoV-2 phylogeny data from December 2019 – January 2022 (above) or December 2019 – September 2025 (below), generated using the Nextstrain ncov GISAID global dataset. Known sites of mutations in 2022 variants of concern or interest are shown in red and then-novel sites predicted based on entropy are shown in orange.
B) Heatmap of the proportion of SARS-CoV-2 spike sequences that contain a mutation at the positions predicted by the WobbRBD 2022 entropy plots by strain. Strains are listed in descending chronological order, with a solid black line denoting the cut-off for variants considered for WobbRBD design. Proportions and sequencing data were collected from CoV-Spectrum, from the World dataset across All Times on October 24, 2025[20].

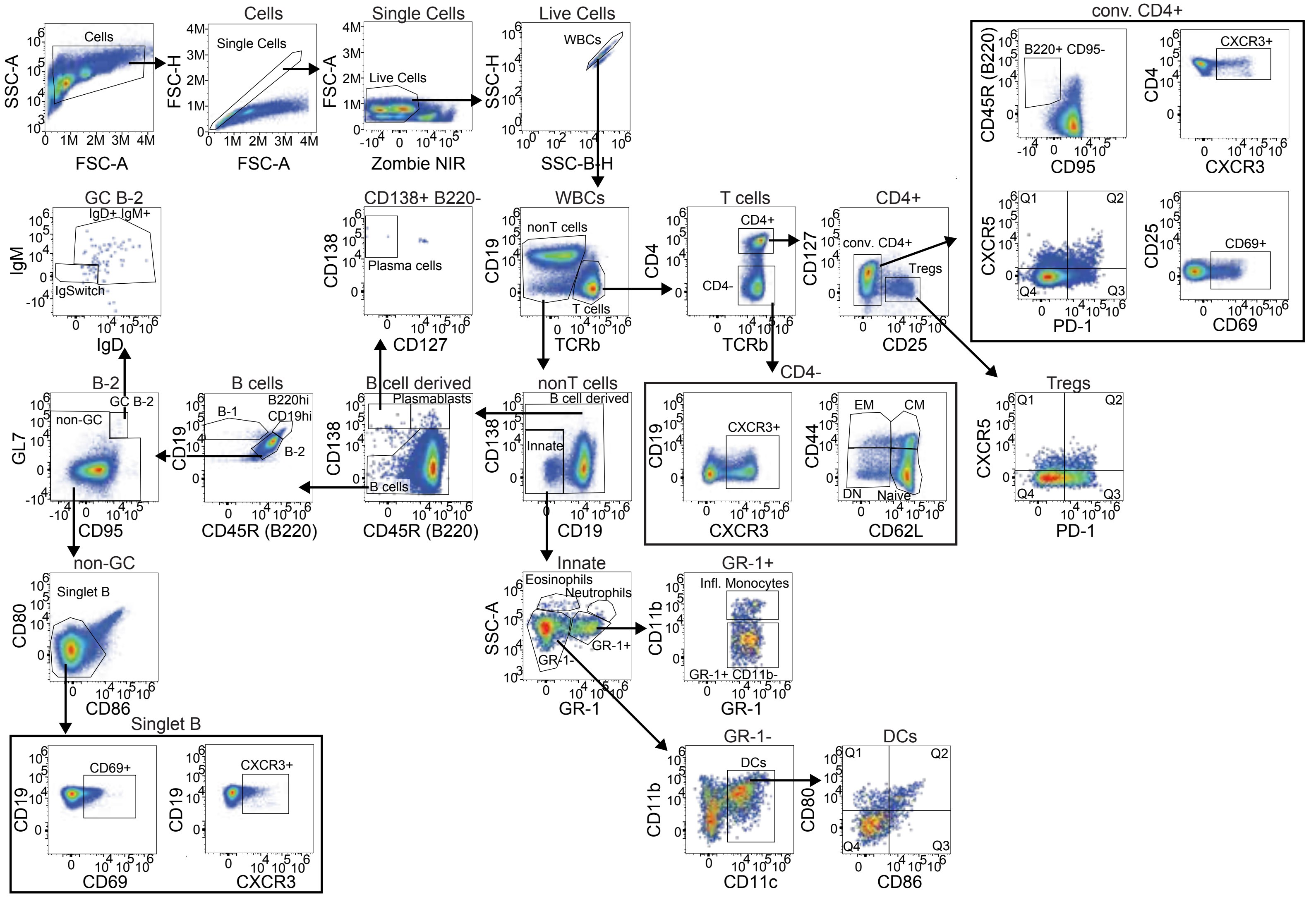

**Supplementary Figure 2. Flow cytometry panel 1 gating scheme (Day 3) –** Gating for the flow panel used for day 3 post-vaccination flow cytometry collection.

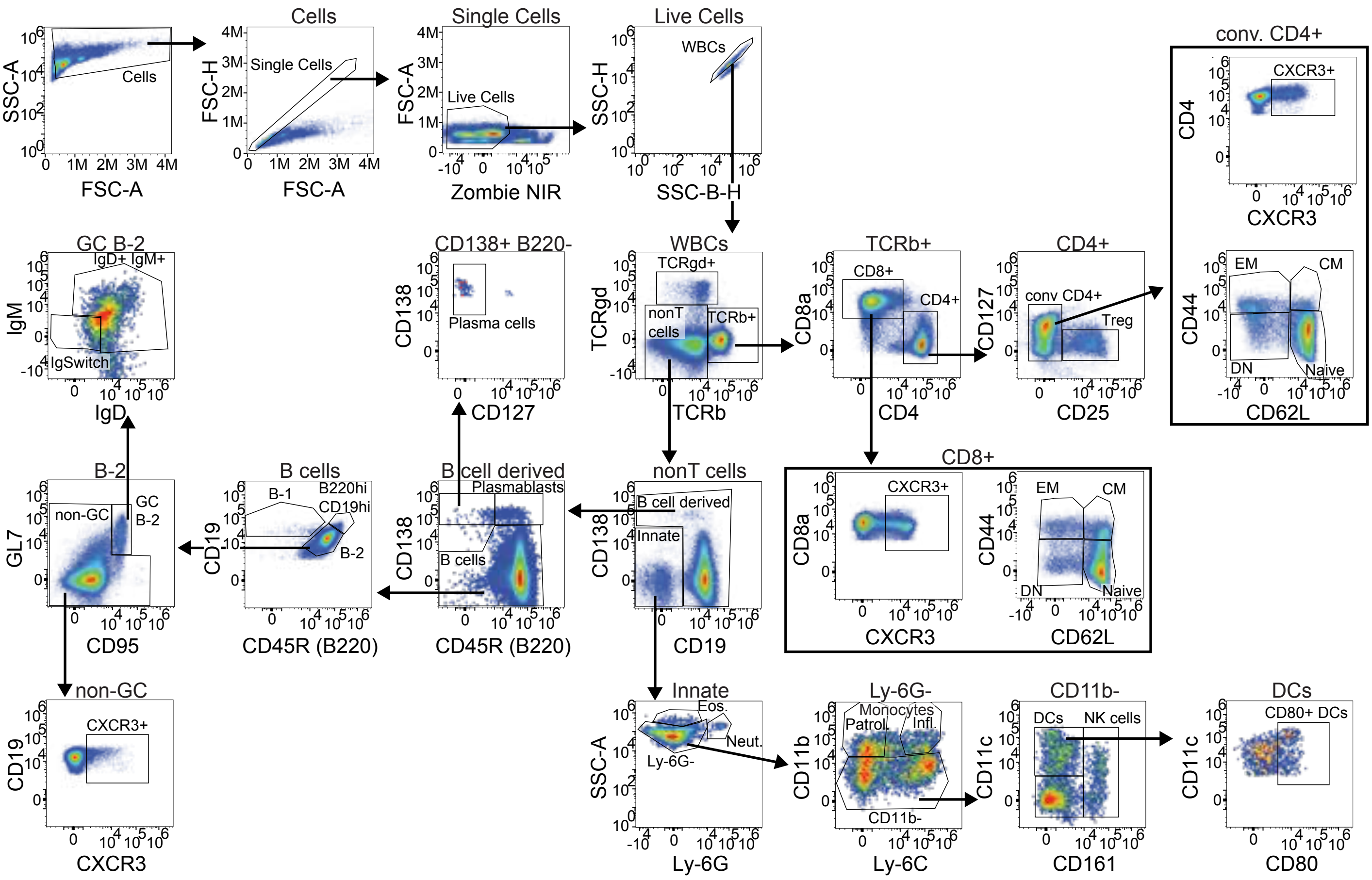

**Supplementary Figure 3. Flow cytometry panel 2 gating scheme (Day 7, 14) –**
Gating for the flow panel used for days 7 and 14 post-vaccination flow cytometry collection.

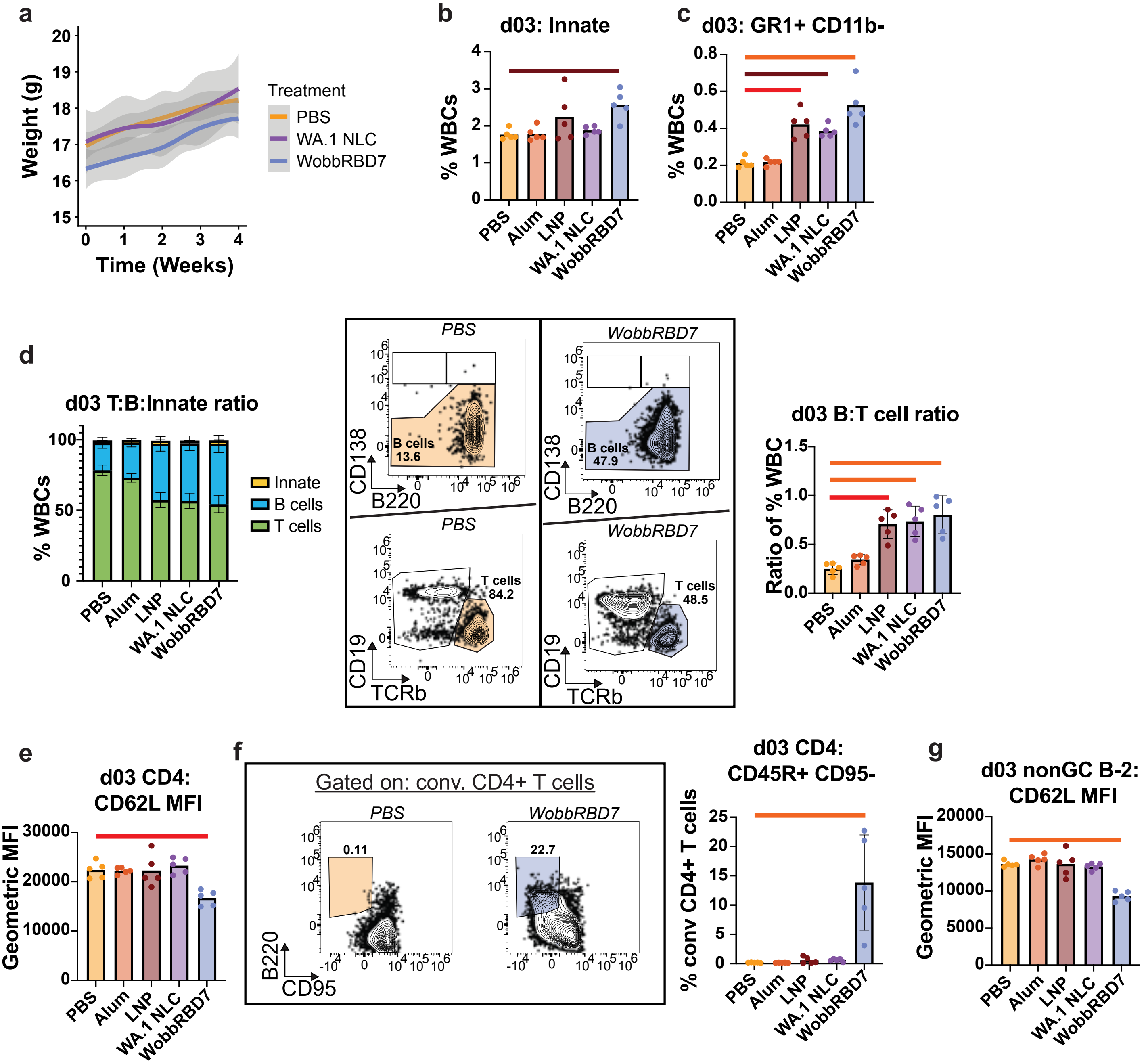

**Supplementary Figure 4. Additional flow cytometry plots for day 3 –**
Additional flow cytometry plots from the experiment described in Figure 3.
A) Weight curve for mice vaccinated with PBS (yellow), 1 µg WA.1 saRNA-NLC (purple), or 1 µg WobbRBD7 saRNA-NLC (blue) over 4 weeks post-administration. Plots were generated using the Loess model from 'geom_smooth' in the R library 'ggplot2.'
B-C, E-G) Additional plots from the experiment described in Figure 3, similar to Fig. 3C-I.
D) Average proportions of white blood cells (WBCs) by treatment. Flow plots show the relevant gates for the PBS (yellow) and WobbRBD7 (blue) with the relevant percent of the WBC gate.

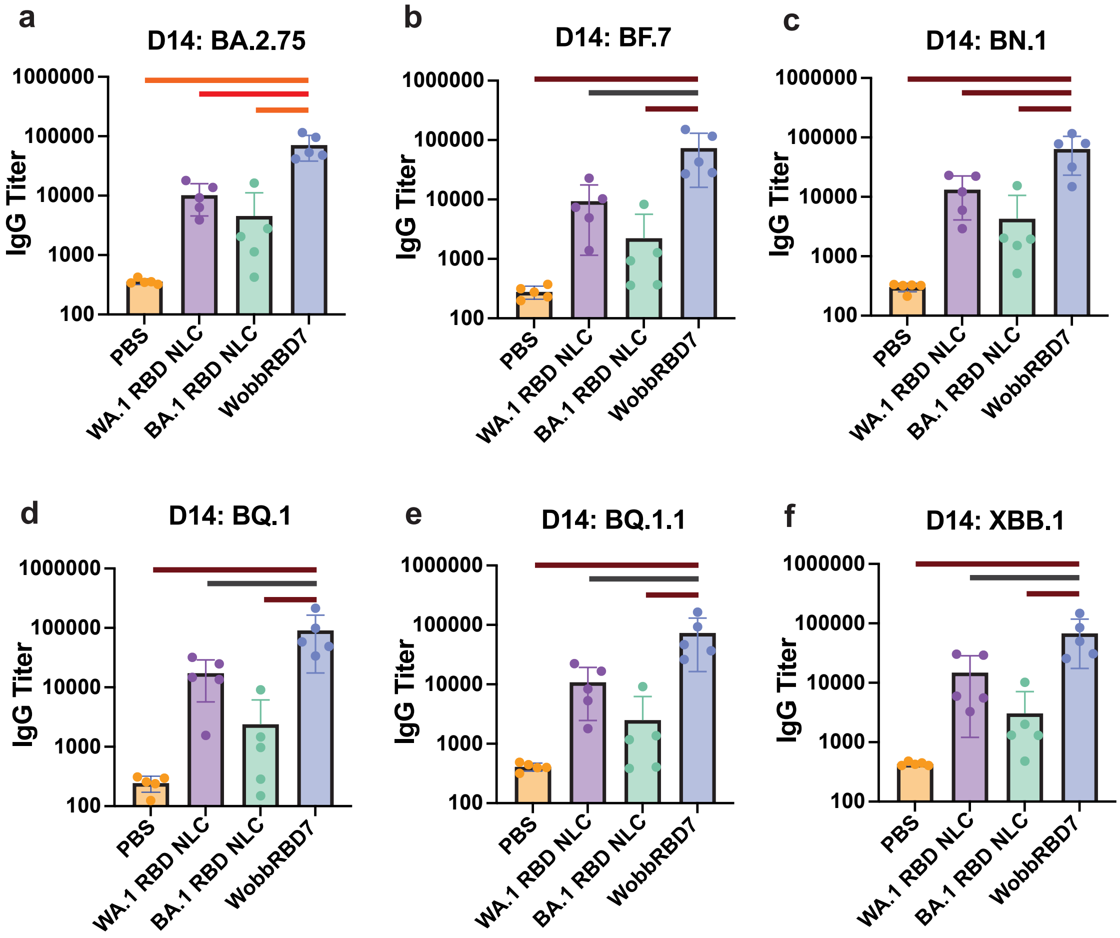

**Supplementary Figure 5. Additional SARS-CoV-2 Spike IgG Titers –**
Additional serum IgG titer plots from the experiment described in Fig. 5C-D.

**Supplementary Table 1. Wobble construct sequences**

A table of DNA sequences for each single strain and wobble construct utilized, with degenerate bases for wobble sites and highlighting to denote wobble regions.

| Name | Encoded protein(s) | Sequence |
|---|---|---|
| sarscov2_wuhan_spike_rbd_signalpep | WA.1 RBD | ATGGAAACCGACACACTGCTGCTGTGGGTGCTGCTCCTGTGGGTCCCCGGCTCCACCGGCGTGCAGCCTACCGAGAGCATCGTGCGGTTCCCCAAC<br>ATCACCAACCTGTGTCCTTTCGGCGAGGTGTTCAATGCCACAAGATTCGCCAGCGTGTACGCTTGGAACCGGAAAAGAATCAGCAATTGCGTGGCCGA<br>CTACAGCGTGCTGTATAACAGCGCCTCTTTTTCTACCTTCAAGTGCTACGGCGTGTCCCCAACCAAGCTGAACGACCTGTGCTTCACCAACGTGTACG<br>CCGACAGCTTCGTGATTAGAGGAGATGAGGTGCGGCAGATCGCTCCTGGCCAGACCGGCAAGATCGCCGATTACAACTATAAGCTGCCTGACGACTT<br>CACCGGATGTGTGATCGCCTGGAATTCTAACAACCTGGACAGCAAGGTGGGCGGCAACTACAACTACCTGTACAGACTGTTCAGAAAGAGCAACCTGA<br>AACCTTTTGAGAGAGATATCTCTACCGAGATCTACCAGGCCGGCAGCACACCCTGCAACGGAGTGGAAGGCTTCAACTGCTACTTCCCTCTGCAGAGC<br>TACGGATTTCAGCCCACAAACGGCGTTGGCTACCAACCATACCGCGTGGTGGTCCTGAGCTTTGAACTGCTGCACGCCCCTGCTACAGTGTGCGGCC<br>CTAAGAAGTCCACAAATCTGGTGAAGAACAAATGTGTGAACTTCAATTTCAACGGCTAA |
| sarscov2_omicron.ba.1_spike_rbd_signalpep | BA.1 RBD | ATGGAAACCGACACACTGCTGCTGTGGGTGCTGCTCCTGTGGGTCCCCGGCTCCACCGGCGTGCAGCCTACCGAGAGCATCGTGCGGTTCCCCAAC<br>ATCACCAACCTGTGTCCTTTCGACGAGGTGTTCAATGCCACAAGATTCGCCAGCGTGTACGCTTGGAACCGGAAAAGAATCAGCAATTGCGTGGCCGA<br>CTACAGCGTGCTGTATAACCTGGCCCCCTTTTTCACCTTCAAGTGCTACGGCGTGTCCCCAACCAAGCTGAACGACCTGTGCTTCACCAACGTGTACG<br>CCGACAGCTTCGTGATTAGAGGAGATGAGGTGCGGCAGATCGCTCCTGGCCAGACCGGCAACATCGCCGATTACAACTATAAGCTGCCTGACGACTTC<br>ACCGGATGTGTGATCGCCTGGAATTCTAACAAGCTGGACAGCAAGGTGAGCGGCAACTACAACTACCTGTACAGACTGTTCAGAAAGAGCAACCTGAA<br>ACCTTTTGAGAGAGATATCTCTACCGAGATCTACCAGGCCGGCAACAAGCCCTGCAACGGAGTGGCCGGCTTCAACTGCTACTTCCCTCTGCGGAGCT<br>ACAGCTTTCGGCCCACATACGGCGTTGGCCACCAACCATACCGCGTGGTGGTCCTGAGCTTTGAACTGCTGCACGCCCCTGCTACAGTGTGCGGCCC<br>TAAGAAGTCCACAAATCTGGTGAAGAACAAATGTGTGAACTTCAATTTCAACGGCTAA |
| sarscov2_flex1_spike_rbd_signalpep | WobbRBD1 | ATGGAAACCGACACACTGCTGCTGTGGGTGCTGCTCCTGTGGGTCCCCGGCTCCACCGGCGTGCAGCCTACCGAGAGCATCGTGCGGTTCCCCAAC<br>ATCACCAACCTGTGTCCTTTCSRYGAGGTGTTCAATGCCACAAVRTTCGCCAGCGTGTACGCTTGGAACCGGAAAAGAATCAGCAATTGCGTGGCCGA<br>CTACAGCGTGCTGTATAACAGCGCCTCTTTTTCTACCTTCAAGTGCTACGGCGTGTCCCCAACCAAGCTGAACGACCTGTGCTTCACCAACGTGTACG<br>CCGACAGCTTCGTGATTAGAGGAGATGAGGTGCGGCAGATCGCTCCTGGCCAGACCGGCAAGATCGCCGATTACAACTATAAGCTGCCTGACGACTT<br>CACCGGATGTGTGATCGCCTGGAATTCTAACAACCTGGACAGCAAGGTGGGCGGCAACTACAACTACCTGTACAGACTGTTCAGAAAGAGCAACCTGA<br>AACCTTTTGAGAGAGATATCTCTACCGAGATCTACCAGGCCGGCAGCACACCCTGCAACGGAGTGGAAGGCTTCAACTGCTACTTCCCTCTGCAGAGC<br>TACGGATTTCAGCCCACAAACGGCGTTGGCTACCAACCATACCGCGTGGTGGTCCTGAGCTTTGAACTGCTGCACGCCCCTGCTACAGTGTGCGGCC<br>CTAAGAAGTCCACAAATCTGGTGAAGAACAAATGTGTGAACTTCAATTTCAACGGCTAA |
| sarscov2_flex2_spike_rbd_signalpep | WobbRBD2 | ATGGAAACCGACACACTGCTGCTGTGGGTGCTGCTCCTGTGGGTCCCCGGCTCCACCGGCGTGCAGCCTACCGAGAGCATCGTGCGGTTCCCCAAC<br>ATCACCAACCTGTGTCCTTTCGGCGAGGTGTTCAATGCCACAAGATTCGCCAGCGTGTACGCTTGGAACCGGAAAAGAATCAGCAATTGCGTGGCCGA<br>CTACAGCGTGCTGTATAACTYNGCCYYRTTTTHYRCNTTCAAGTGCTACGGCGTGTCCCCAACCAAGCTGAACGACCTGTGCTTCACCAACGTGTACGC<br>CGACAGCTTCGTGATTAGAGGAGATGAGGTGCGGCAGATCGCTCCTGGCCAGACCGGCAAGATCGCCGATTACAACTATAAGCTGCCTGACGACTTCA<br>CCGGATGTGTGATCGCCTGGAATTCTAACAACCTGGACAGCAAGGTGGGCGGCAACTACAACTACCTGTACAGACTGTTCAGAAAGAGCAACCTGAAA<br>CCTTTTGAGAGAGATATCTCTACCGAGATCTACCAGGCCGGCAGCACACCCTGCAACGGAGTGGAAGGCTTCAACTGCTACTTCCCTCTGCAGAGCTA<br>CGGATTTCAGCCCACAAACGGCGTTGGCTACCAACCATACCGCGTGGTGGTCCTGAGCTTTGAACTGCTGCACGCCCCTGCTACAGTGTGCGGCCCT<br>AAGAAGTCCACAAATCTGGTGAAGAACAAATGTGTGAACTTCAATTTCAACGGCTAA |
| sarscov2_flex3_spike_rbd_signalpep | WobbRBD3 | ATGGAAACCGACACACTGCTGCTGTGGGTGCTGCTCCTGTGGGTCCCCGGCTCCACCGGCGTGCAGCCTACCGAGAGCATCGTGCGGTTCCCCAAC<br>ATCACCAACCTGTGTCCTTTCGGCGAGGTGTTCAATGCCACAAGATTCGCCAGCGTGTACGCTTGGAACCGGAAAAGAATCAGCAATTGCGTGGCCGA<br>CTACAGCGTGCTGTATAACAGCGCCTCTTTTTCTACCTTCAAGTGCTACGGCGTGTCCCCAACCAAGCTGAACGACCTGTGCTTCACCAACGTGTACG<br>CCGACAGCTTCGTGATTAGAGGARAYGAGGTGAGNCAGATCGCTCCTGGCCAGACCGGCANAATCGCCGATTACAACTATAAGCTGCCTGACGACTTC<br>ACCGGATGTGTGATCGCCTGGAATTCTAACAACCTGGACAGCAAGGTGGGCGGCAACTACAACTACCTGTACAGACTGTTCAGAAAGAGCAACCTGAA<br>ACCTTTTGAGAGAGATATCTCTACCGAGATCTACCAGGCCGGCAGCACACCCTGCAACGGAGTGGAAGGCTTCAACTGCTACTTCCCTCTGCAGAGCT<br>ACGGATTTCAGCCCACAAACGGCGTTGGCTACCAACCATACCGCGTGGTGGTCCTGAGCTTTGAACTGCTGCACGCCCCTGCTACAGTGTGCGGCCC<br>TAAGAAGTCCACAAATCTGGTGAAGAACAAATGTGTGAACTTCAATTTCAACGGCTAA |
| sarscov2_flex4_spike_rbd_signalpep | WobbRBD4 | ATGGAAACCGACACACTGCTGCTGTGGGTGCTGCTCCTGTGGGTCCCCGGCTCCACCGGCGTGCAGCCTACCGAGAGCATCGTGCGGTTCCCCAAC<br>ATCACCAACCTGTGTCCTTTCGGCGAGGTGTTCAATGCCACAAGATTCGCCAGCGTGTACGCTTGGAACCGGAAAAGAATCAGCAATTGCGTGGCCGA<br>CTACAGCGTGCTGTATAACAGCGCCTCTTTTTCTACCTTCAAGTGCTACGGCGTGTCCCCAACCAAGCTGAACGACCTGTGCTTCACCAACGTGTACG<br>CCGACAGCTTCGTGATTAGAGGAGATGAGGTGCGGCAGATCGCTCCTGGCCAGACCGGCAAGATCGCCGATTACAACTATAAGCTGCCTGACGACTT<br>CACCGGATGTGTGATCGCCTGGAATTCTAANAANCTGGACAGCAAGGTGGRYGGCAACYAYAACTACCDRTACAGACTGTTCAGAAAGAGCAACCTGA<br>AACCTTTTGAGAGAGATATCTCTACCGAGATCTACCAGGCCGGCAGCACACCCTGCAACGGAGTGGAAGGCTTCAACTGCTACTTCCCTCTGCAGAGC<br>TACGGATTTCAGCCCACAAACGGCGTTGGCTACCAACCATACCGCGTGGTGGTCCTGAGCTTTGAACTGCTGCACGCCCCTGCTACAGTGTGCGGCC<br>CTAAGAAGTCCACAAATCTGGTGAAGAACAAATGTGTGAACTTCAATTTCAACGGCTAA |
| sarscov2_flex5_spike_rbd_signalpep | WobbRBD5 | ATGGAAACCGACACACTGCTGCTGTGGGTGCTGCTCCTGTGGGTCCCCGGCTCCACCGGCGTGCAGCCTACCGAGAGCATCGTGCGGTTCCCCAAC<br>ATCACCAACCTGTGTCCTTTCGGCGAGGTGTTCAATGCCACAAGATTCGCCAGCGTGTACGCTTGGAACCGGAAAAGAATCAGCAATTGCGTGGCCGA<br>CTACAGCGTGCTGTATAACAGCGCCTCTTTTTCTACCTTCAAGTGCTACGGCGTGTCCCCAACCAAGCTGAACGACCTGTGCTTCACCAACGTGTACG<br>CCGACAGCTTCGTGATTAGAGGAGATGAGGTGCGGCAGATCGCTCCTGGCCAGACCGGCAAGATCGCCGATTACAACTATAAGCTGCCTGACGACTT<br>CACCGGATGTGTGATCGCCTGGAATTCTAACAACCTGGACAGCAAGGTGGGCGGCAACTACAACTACCTGTACAGACTGTTCAGAAAGAGCAACCTGA<br>AACCTTTTGAGAGAGATATCTCTACCGAGATCTACCAGGCCGGCADYANACCCTGCAANGGAGTGVARGGCTTCAACTGCTACTTCCCTCTGCAGAGC<br>TACGGATTTCAGCCCACAAACGGCGTTGGCTACCAACCATACCGCGTGGTGGTCCTGAGCTTTGAACTGCTGCACGCCCCTGCTACAGTGTGCGGCC<br>CTAAGAAGTCCACAAATCTGGTGAAGAACAAATGTGTGAACTTCAATTTCAACGGCTAA |
| sarscov2_flex6_spike_rbd_signalpep | WobbRBD6 | ATGGAAACCGACACACTGCTGCTGTGGGTGCTGCTCCTGTGGGTCCCCGGCTCCACCGGCGTGCAGCCTACCGAGAGCATCGTGCGGTTCCCCAAC<br>ATCACCAACCTGTGTCCTTTCGGCGAGGTGTTCAATGCCACAAGATTCGCCAGCGTGTACGCTTGGAACCGGAAAAGAATCAGCAATTGCGTGGCCGA<br>CTACAGCGTGCTGTATAACAGCGCCTCTTTTTCTACCTTCAAGTGCTACGGCGTGTCCCCAACCAAGCTGAACGACCTGTGCTTCACCAACGTGTACG<br>CCGACAGCTTCGTGATTAGAGGAGATGAGGTGCGGCAGATCGCTCCTGGCCAGACCGGCAAGATCGCCGATTACAACTATAAGCTGCCTGACGACTT<br>CACCGGATGTGTGATCGCCTGGAATTCTAACAACCTGGACAGCAAGGTGGGCGGCAACTACAACTACCTGTACAGACTGTTCAGAAAGAGCAACCTGA<br>AACCTTTTGAGAGAGATATCTCTACCGAGATCTACCAGGCCGGCAGCACACCCTGCAACGGAGTGGAAGGCTTCAACTGCTACTTCCCTCTGCWRAGC<br>TACRGYTTTCRRCCCACAWAYGGCGTTGGCYAYCAACCATACCGCGTGGTGGTCCTGAGCTTTGAACTGCTGCACGCCCCTGCTACAGTGTGCGGCC<br>CTAAGAAGTCCACAAATCTGGTGAAGAACAAATGTGTGAACTTCAATTTCAACGGCTAA |
| sarscov2_fullflex_spike_rbd_signalpep | WobbRBD7 | ATGGAAACCGACACACTGCTGCTGTGGGTGCTGCTCCTGTGGGTCCCCGGCTCCACCGGCGTGCAGCCTACCGAGAGCATCGTGCGGTTCCCCAAC<br>ATCACCAACCTGTGTCCTTTCSRYGAGGTGTTCAATGCCACAAVRTTCGCCAGCGTGTACGCTTGGAACCGGAAAAGAATCAGCAATTGCGTGGCCGA<br>CTACAGCGTGCTGTATAACTYNGCCYYRTTTTHYRCNTTCAAGTGCTACGGCGTGTCCCCAACCAAGCTGAACGACCTGTGCTTCACCAACGTGTACGC<br>CGACAGCTTCGTGATTAGAGGARAYGAGGTGAGNCAGATCGCTCCTGGCCAGACCGGCANAATCGCCGATTACAACTATAAGCTGCCTGACGACTTCA<br>CCGGATGTGTGATCGCCTGGAATTCTAANAANCTGGACAGCAAGGTGGRYGGCAACYAYAACTACCDRTACAGACTGTTCAGAAAGAGCAACCTGAAA<br>CCTTTTGAGAGAGATATCTCTACCGAGATCTACCAGGCCGGCADYANACCCTGCAANGGAGTGVARGGCTTCAACTGCTACTTCCCTCTGCWRAGCTA<br>CRGYTTTCRRCCCACAWAYGGCGTTGGCYAYCAACCATACCGCGTGGTGGTCCTGAGCTTTGAACTGCTGCACGCCCCTGCTACAGTGTGCGGCCCT<br>AAGAAGTCCACAAATCTGGTGAAGAACAAATGTGTGAACTTCAATTTCAACGGCTAA |

**Supplementary Table 2. saRNA-NLC quality control metrics**

A table of quality control metrics for each saRNA-NLC utilized.

| Name | Z-average (DLS; nm) | Particle Size Distribution Spread (DLS) | RNA concentration (Ribogreen; mg/ml) | RNA Main Band % Intensity (Gel Elecrophoresis) | Rnase Protection (Rnase:RNA 1:200; Gel Electrophoresis) |
|---|---|---|---|---|---|
| WA.1 saRNA-NLC | 67.0 ± 0.2 | 0.20 ± 0.02 | 0.046 | 64% | 94% |
| Omicron saRNA-NLC | 65 ± 1 | 0.18 ± 0.01 | 0.043 | 65% | 85% |
| WobbRBD1 saRNA-NLC | 64.0 ± 0.9 | 0.17 ± 0.00 | 0.049 | 65% | 76% |
| WobbRBD2 saRNA-NLC | 64 ± 1 | 0.16 ± 0.02 | 0.045 | 66% | 81% |
| WobbRBD3 saRNA-NLC | 67 ± 1 | 0.18 ± 0.01 | 0.05 | 71% | 78% |
| WobbRBD4 saRNA-NLC | 66 ± 1 | 0.17 ± 0.02 | 0.051 | 75% | 93% |
| WobbRBD5 saRNA-NLC | 65 ± 1 | 0.18 ± 0.02 | 0.05 | 71% | 90% |
| WobbRBD6 saRNA-NLC | 64 ± 1 | 0.17 ± 0.01 | 0.047 | 76% | 84% |
| WobbRBD7 saRNA-NLC | 75 ± 1 | 0.20 ± 0.01 | 0.056 | 80% | N/A |

**Key Resources Table**

| Category | Reagent or Resource | Source | Identifier (Cat#, clone) | Flow Cytometry Panel 1 (B cell stim, Day 3) | Flow Cytometry Panel 2 (CBC+, Day 7+14) |
|---|---|---|---|---|---|
| Flow cytometry antibodies | CXCR3-BV421 | Biolegend | 126521, CXCR3-173 | Y | Y |
| | CD62L-BUV737 | Thermo | 367-0621-80, MEL-14 | Y | Y |
| | CD21/35-RealBlue780 | BD Biosciences | 755428, 7G6 | Y | Y |
| | IgD-PE-Cy5.5 | Elabscience | E-AB-F1189, 11-26.2a | Y | Y |
| | CD80-BV786 | BD Biosciences | 740888, 16-10A1 | Y | Y |
| | CD11b-APC-Fire810 | Biolegend | 101288, M1/70 | Y | Y |
| | CD86-Spark Plus B550 | Biolegend | 105061, GL-1 | Y | N |
| | CD44-AlexaFluor700 | Biolegend | 103025, IM7 | Y | Y |
| | PD-1-PE-Cy7 | Biolegend | 135215, 29F.1A12 | Y | N |
| | CD19-BUV615 | Thermo | 366-0193-82, eBio1D3 (1D3) | Y | N |
| | CD4-BV480 | Thermo | 414-0042-82, RM4-5 | Y | Y |
| | GR-1-BV570 | Biolegend | 108431, RB6-8C5 | Y | N |
| | CD95-BUV496 | BD Biosciences | 741103, JO2 | Y | Y |
| | CD69-BUV395 | BD Biosciences | 569367, H1.2F3 | Y | N |
| | CD45R (B220)- BV510 | Biolegend | 103248, RA3-6B2 | Y | N |
| | CD25-BV650 | Biolegend | 102037, PC61 | Y | Y |
| | CD138-BV750 | BD Biosciences | 747070, 281-2 | Y | Y |
| | CXCR5-PE-Dazzle594 | Biolegend | 145521, L138D7 | Y | Y |
| | CD11c-BUV563 | Thermo | 365-0114-82, N418 | Y | Y |
| | GL7-PE | BD Biosciences | 561530, GL7 | Y | Y |
| | TCR-beta-BUV805 | BD Biosciences | 748405, H57-597 | Y | Y |
| | IgM-APC-Fire750 | Biolegend | 406538, RMM-1 | Y | Y |
| | CD127-BUV661 | Thermo | 376-1271-82, A7R34 | Y | N |
| | CCR4-PE-Cy7 | Biolegend | 131213, 2G12 | N | Y |
| | CD8a-BUV615 | Thermo | 613004, 53-6.7 | N | Y |
| | CD19-eFluor450 | Thermo | 48-0193-82, eBio1D3 (1D3) | N | Y |
| | PD-L1-BV605 | BD Biosciences | 568559, 10F.9G2 | N | Y |
| | CD45R (B220)- SparkNIR685 | Biolegend | 103268, RA3-6B2 | N | Y |
| | Ly6C-BV570 | Biolegend | 128030, HK1.4 | N | Y |
| | Ly6G-SparkBlue550 | Biolegend | 127664, 1A8 | N | Y |
| | CD161-BV711 | Biolegend | 108745, PK136 | N | Y |
| | TCRg/d-PE-Cy5 | Thermo | 15-5711-82, eBioGL3 (GL-3 GL3) | N | Y |
| | PD-L2-BUV395 | BD Biosciences | 565102, Ty25 | N | Y |
| | Zombie NIR | Biolegend | 423106 | Y | Y |
| Flow cytometry reagents | Brilliant Stain Buffer Plus | BD Biosciences | 566385 | | |
| | TruStain FcX (anti-mouse CD16/32) antibody | Biolegend | 101320 | | |
| General reagents | Phosphate Buffered Saline (PBS) | Corning | 21-040-CV | | |
| | Fetal Bovine Serum (FBS), Heat Inactivated | Corning | 35-011-CV | | |
| | Ethylenediamine Tetraacetate Acid (EDTA) | Fisher Bioreagents | BP2482-100 | | |
| | Goldenrod Animal Lancet 5mm | Braintree | GR 5MM | | |
| | Alhydrogel 2% | InvivoGen | 5808-46-02 | | |
| | Eagle's Minimum Essential Medium with 1.5 g/L sodium bicarbonate, non-essential amino acids, L-glutamine and sodium pyruvate | Corning | 10-009-CV | | |
| | U-100 Insulin syringes 0.3ml 29G x 1/2" | Exel International | 26018 | | |
| | 96 well V-bottom assay plate | Costar | 3897 | | |
| | 1.5ml pestles | VWR | 64788-10 | | |
| | HEp-2 cells | ATCC | CCL-23 | | |
| | Vivaspin 2 10 kDa MWCO concentrator | Cytiva | 28932247 | | |
| | Human SARS-CoV-2 RBD ELISA kit | Invitrogen | EH492RB | | |
| | V-PLEX SARS-CoV-2 Panel 34 (Mouse IgG) kit | MesoScale Discovery | K15694U | | |
